# An introduction to MPEG-G, the new ISO standard for genomic information representation

**DOI:** 10.1101/426353

**Authors:** Claudio Albert, Tom Paridaens, Jan Voges, Daniel Naro, Junaid J. Ahmad, Massimo Ravasi, Daniele Renzi, Giorgio Zoia, Paolo Ribeca, Idoia Ochoa, Marco Mattavelli, Jaime Delgado, Mikel Hernaez

## Abstract

The MPEG-G standardization initiative is a coordinated international effort to specify a compressed data format that enables large scale genomic data to be processed, transported and shared. The standard consists of a set of specifications (i.e., a book) describing: i) a normative format syntax, and ii) a normative decoding process to retrieve the information coded in a compliant file or bitstream. Such decoding process enables the use of leading-edge compression technologies that have exhibited significant compression gains over currently used formats for storage of unaligned and aligned sequencing reads. Additionally, the standard provides a wealth of much needed functionality, such as selective access, data aggregation, application programming interfaces to the compressed data, standard interfaces to support data protection mechanisms, support for streaming and a procedure to assess the conformance of implementations. ISO/IEC is engaged in supporting the maintenance and availability of the standard specification, which guarantees the perenniality of applications using MPEG-G. Finally, the standard ensures interoperability and integration with existing genomic information processing pipelines by providing support for conversion from the FASTQ/SAM/BAM file formats.

In this paper we provide an overview of the MPEG-G specification, with particular focus on the main advantages and novel functionality it offers. As the standard only specifies the decoding process, encoding performance, both in terms of speed and compression ratio, can vary depending on specific encoder implementations, and will likely improve during the lifetime of MPEG-G. Hence, the performance statistics provided here are only indicative baseline examples of the technologies included in the standard.

## INTRODUCTION

The development and rapid progress of High-Throughput Sequencing (HTS) technologies holds the potential to enable the use of genomic information as an everyday practice in a number of fields. With the release of the latest generation of Illumina HTS machines (HiSeq X and NovaSeq series), the cost of sequencing a whole human genome has dropped to less than US$1,000. In the next few years such cost is expected to progressively drop further, to about US$100. Today, a single sequencing system can deliver the equivalent of 10,000 whole human genomes sequenced per year, which translates to more than 1 PB of data. This fact has lead to the forecast that the amount of generated genomic data will soon surpass the volume of astronomical data [1]. By that point the IT costs associated to storing, transmitting and processing the large volumes of genomic data will largely exceed the costs of sequencing. In addition, the lack of an appropriate compressed data representation is widely recognized as a critical element limiting the potential for genomic data to be used in a wide range of scientific and public health scenarios [2]. Note that this is not due to a lack of specialized compressors for genomic data *per se* (see [3] and references therein), but rather to the absence of efficient, perennial and reliable solutions able to offer a complete framework—beyond compression—for the representation of genomic information.

Motivated by these facts, the Moving Picture Experts Group (MPEG)—a joint working group of the International Standardization Organization (ISO) and the International Electrotechnical Commission (IEC)—is working with ISO Technical Committee 276/Working Group 5, integrators of biological data workflows, to produce MPEG-G, a new open standard to compress, store, transmit and process sequencing data. In its 30 years of activity, MPEG has already developed several generations of successful standards that have transformed the world of media from analog to digital (e.g., MP3 and AAC for audio, and AVC/H.264 and HEVC/H.265 for video [4]). These standards enabled the interoperability and integration we all can witness today in the field of digital media.

MPEG-G has been developed following the open and rigorous process adopted by MPEG for all of its standards. The first step was the production of a list of requirements for the compressed representation of raw and aligned reads produced during primary and secondary data analysis. The standard also includes requirements for the efficient transport of, and selective access to, compressed genomic data. The process of identifying all requirements was a wide interdisciplinary effort sustained by experts from different domains including bioinformatics, biology, information theory, telecommunication, video and data compression, data storage, and information security [^1^ The identified requirements that were the baseline for the development of the MPEG-G standard are available in full detail in the public document N16323 (MPEG)/N97 (ISO TC276/WG5) [5]].

A Call for Proposals was then issued in June 2016 and 15 responses were received from 17 organizations in October 2016. The identified technologies were evaluated using a number of criteria, including—but not limited to—compression performance, selective access capabilities, and flexibility for efficient coding of a wide variety of sequencing data. Separate assessments for different types of genomic data were performed: sequence reads, quality values, read identifiers, alignment information, and metadata. In addition, a preliminary evaluation of the computational complexity was assessed by measuring encoding and decoding speed as well as memory usage. This ensured that the candidate technologies were compatible with efficient implementations. The support for non-sequential access, extended nucleotide alphabets, encoding of additional metadata (extensibility), and quantized coding of sequencing quality values (often referred to as quality scores) was also considered when evaluating and ranking the submitted proposals.

The most valuable technologies were integrated to provide i) the compression of raw genomic data generated by sequencing technologies, ii) the compression of alignment information associated with genomic data when the latter is considered in the context of a reference sequence, and iii) the definition of a Genomic Information Transport Layer that supports storage and transport. In addition, the MPEG-G standard supports features associated to complex use cases, most of which are not supported by currently existing formats (e.g., FASTQ and SAM/BAM). Notable use cases addressed by MPEG-G include:

- Selective access to compressed data (according to several criteria)
- Data streaming
- Concatenation of compressed files and aggregation of genomic studies
- Enforcement of privacy rules
- Selective encryption of sequencing data and metadata
- Annotation and linkage of genomic segments
- Incremental update of sequencing data and metadata.

Finally, interoperability and integration with existing genomic information processing pipelines is ensured by supporting conversion from/to file formats such as FASTQ and SAM/BAM.

In the following we describe the MPEG-G standard in more detail, with an emphasis on its features and capabilities, and provide a discussion on the role of the standard in the future of genomic data storage, access, sharing, and processing. It should be clarified that the MPEG-G standard only specifies the decoder syntax and the algorithms necessary to extract information from files and streams encoded using MPEG-G. This approach allows for the continuous development of new and more optimized encoders, while maintaining compatibility with any existing standard-compliant decoder.

## RESULTS

### Genomic information representation

MPEG-G technology provides storage and transport capabilities for both raw genomic sequences and genomic sequences aligned to reference genomes. It further supports the representation of both single reference genomes (assemblies) and collections thereof. The representation of genomic sequencing data in MPEG-G is based on the concept of *Genomic Records*. The Genomic Record is a data structure consisting of either a single sequence read or a set of paired sequence reads. If available, it contains associated sequencing and alignment information, a set of read identifiers, and a set of quality scores.

Without breaking traditional approaches, the Genomic Record data structure provides a more compact, simpler and manageable data structure grouping all the information related to a single DNA template: from simple raw sequencing data to sophisticated alignment information. However, even if the Genomic Record is an appropriate data structure for interaction and manipulation of genomic information, it is not a suitable atomic data structure for compression. As an example, when dealing with selective data access, the Genomic Record is a too small unit to allow efficient and fast information retrieval while at the same time being highly compressible.

To facilitate both objectives, Genomic Records are classified and grouped into six *Data Classes* that are defined according to the result of their alignment against one or more reference sequences (e.g., perfect matches in Data Class P, matches containing substitutions only in Data Class M, matches containing indels in Data Class I, and Data Class U containing either reads that could not be mapped or raw sequencing data). To further improve compression efficiency, the information contained in the clustered Genomic Records is split across so-called *Descriptor Streams*. Each Descriptor Stream contains information of a specific type. Examples of these Descriptors Streams are: mapping positions, number of substitutions, and read lengths (see Methods).

The classification of sequence reads into Data Classes enables the development of powerful selective data access mechanisms. To make this possible, MPEG-G introduces the concept of an Access Unit, which is the fundamental structure for coding of and access to information in the compressed domain. Access Units are units of coded genomic information that can be independently accessed and inspected. In fact, Access Units are composed solely of Genomic Records pertaining to a specific Data Class, and thus constitute a data structure capable of providing powerful filtering capabilities for the efficient support of many different use cases. An illustration of the essential elements of Access Units in the MPEG-G *File Format* is shown in Figure 1.

**Figure 1:**
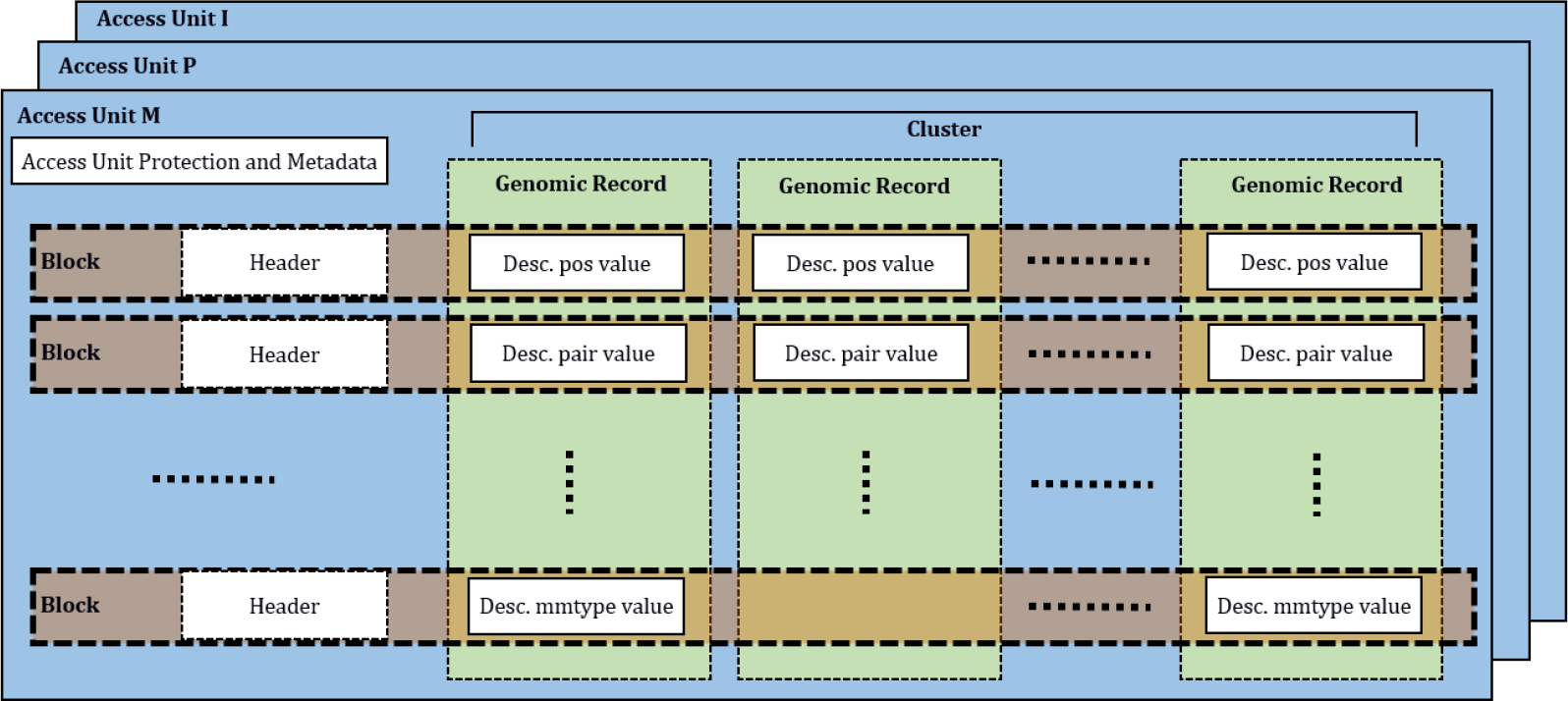
Key elements of an Access Unit in the MPEG-G File Format. Each Access Unit contains Genomic Records belonging to only one Data Class.

Access Units comprise a header and a set of data *Blocks*. The Access Unit header contains the metadata describing the genomic data encoded in the Blocks, such as data type, read count, genomic region the reads are mapped to, presence of multiple alignments, presence of spliced reads, number of coded reads containing substitutions above/below a given threshold, and subsequences (e.g., barcodes from single-cell RNA sequencing experiments), among others. The Blocks contain the coded (i.e., compressed) genomic data. Optionally, additional data structures can be associated to an Access Unit. These data structures can for example contain SAM auxiliary fields or metadata related to protection mechanisms which govern the access to the Access Unit.

To facilitate the storage and transport of genomic information, MPEG-G specifies a digital container for the genomic data, the MPEG-G File Format (Figure 2). As illustrated in the figure, an MPEG-G file is organized in a file header and one or more containers named *Dataset Groups*. Each Dataset Group, at the same time, encapsulates one or more *Datasets*, a header and optional metadata associated to the Dataset Group. Finally, each Dataset contains a header, optional metadata containers and carries one or more Access Units. The nested nature of the File Format allows for efficient queries on and selective access to the compressed data.

**Figure 2:**
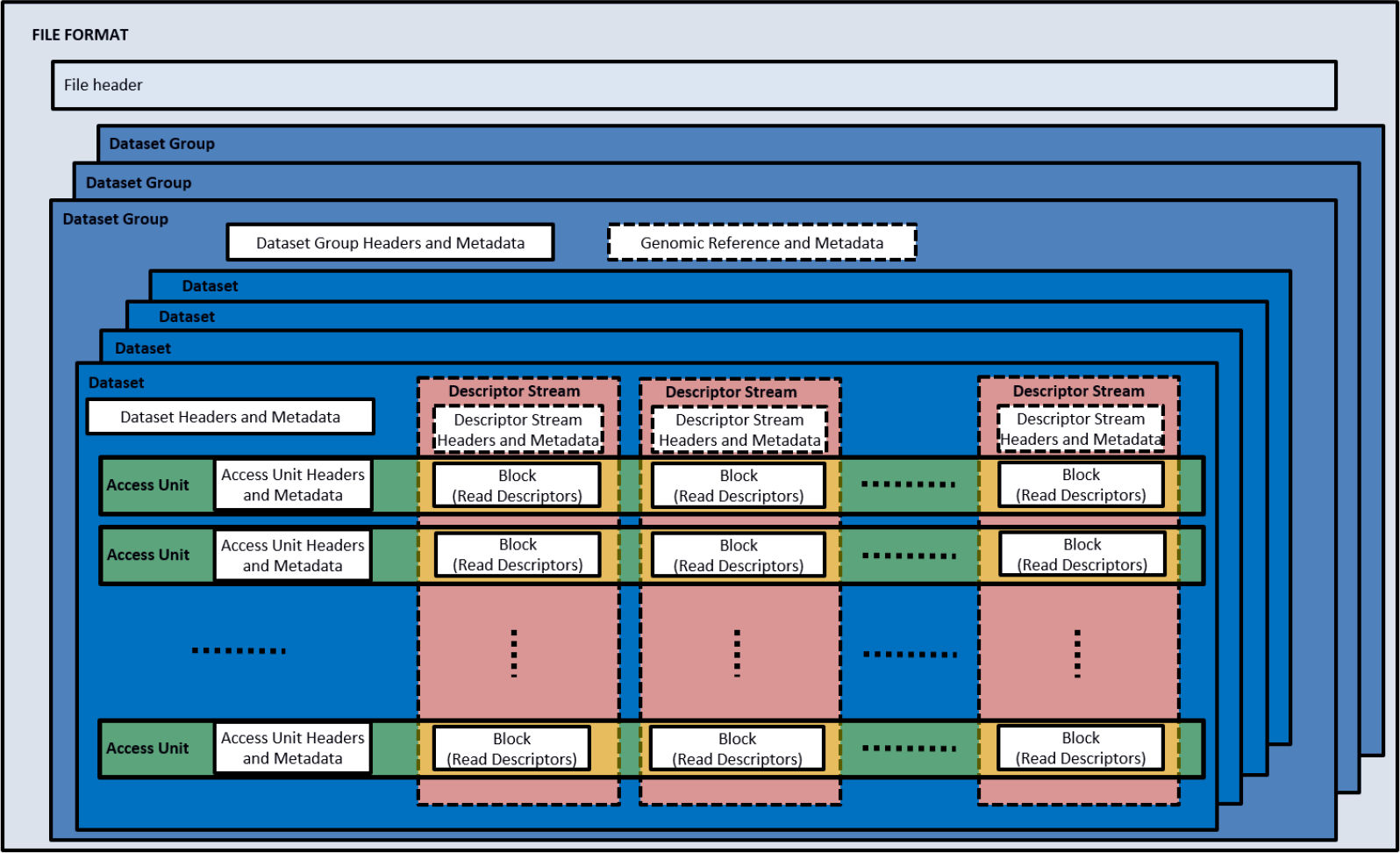
Key elements of the MPEG-G File Format. Multiple Dataset Groups contain multiple Datasets of sequencing data. Each Dataset is composed of Access Units containing Genomic Records pertaining to one specific Data Class. Each Access Unit is composed by Blocks of Read Descriptors.

For example, one could use an MPEG-G file to structure the storage of the genome sequencing data of a trio of individuals (father, mother, child) as follows: there would be three distinct Dataset Groups, one for each individual in the trio. Then, each Dataset Group would contain Datasets related to sequencing runs for the same individual performed either at different moments in time or from different libraries. This example shows how MPEG-G files enable the possibility to encapsulate the entire genomic history of one or more individuals in a unique file including any metadata related to the study, samples, etc.

### Compression capabilities

Sequencing data and their associated metadata are sets of heterogeneous data, each characterized by their own statistical behaviors. Therefore, MPEG-G provides several strategies for classification of these data and their representation. Within MPEG-G, the optimization space for compression performance and selective data access is wide and admits many different solutions, which can be optimized for different applications and even for specific sequencing technologies and species. For example, the data compression mode can be optimized for high compression and indexing (archival), or low latency (streaming applications). Furthermore, aligned reads can be compressed either reference-free or reference-based. In the latter case, MPEG-G supports the use of reference sequences stored in both FASTA and MPEG-G file formats. The used reference sequences can be embedded as Datasets within the same MPEG-G file or stored as external reference sequences, using an unambiguous specification of these external reference sequences. Quality scores can also be compressed either lossless or quantized.

During the development of the MPEG-G specification, the best-performing compression tech-nologies, according to the results of the Call for Proposals, were selected for integration into the MPEG-G specification (see the Methods section for a list of some of these technologies). Note, however, that as a standard development approach, only the decoding process is normative and specified. This guarantees the interoperability of applications implementing the standard, while the encoding process is open to algorithmic and implementation-specific innovations. As such, the compression performance achievable by MPEG-G will vary from encoder to encoder, depending on the individual implementations, and most likely will improve across time. Nevertheless, to give the reader a sense of the compression capabilities achievable by MPEG-G, next we show the compression performance of some of the technologies supported by the standard (and hence implementable in an encoder), and how they compare with current formats [^2^ The specific numbers are taken from the corresponding publications.]. It should be noted that making these technologies MPEG-G compatible may require some syntax modifications to produce MPEG-G compatible descriptor streams, possible leading to slightly different compression performances. In addition, since an MPEG-G file provides additional functionality beyond pure compression, some minor overhead is to be expected.

In todays common practice, SAM files are often stored or transmitted in the form of BAM files, which are essentially block-wise binarized and gzipped SAM files. The CRAM format [6], which efficiently represents aligned data, is also seeing wide acceptance. On the contrary, FASTQ files are generally compressed with a general lossless compressor (e.g., gzip).

For example, the information contained in the FASTQ file (i.e., identifiers, reads, and quality scores) could be compressed by an MPEG-G encoder in a similar way as done in FaStore [7], one of the best compressors for FASTQ files presented in the literature. In this case, whereas gzip reduces the FASTQ file by more than 70% on average, FaStore achieves more than 85% reduction (for all supported modes, including lossless). When it comes to aligned data, an MPEG-G encoder could use a compression method comparable to that of DeeZ [8], which is able to compress a 437 GB H. Sapiens SAM file to about 63 GB, as compared to 75 GB by CRAM (Scramble) or 106 GB by BAM [8]. Regarding quantization of quality values, methods like QVZ [9] and CALQ [10] could be applied yielding overall compression gains of 10x over BAM, while preserving, or even improving, variant calling performance [11].

Finally, technology employed for entropy coding (see Methods) has been shown to compress the MPEG-G descriptor stream test set down to about 21% of the uncompressed size at an average processing speed of more than 25 MiB/s per descriptor stream (with single-threaded encoding) [12].

As discussed above, these are only examples of a possible coding performance achievable, and compression ratios as well as compression speeds may vary according to the specific statistical characteristics of each data set and according to the quality and optimization capabilities of the encoder. Note also that some minor overhead is expected to be incurred due to the encapsulation of each Dataset into an MPEG-G compatible file.

### MPEG-G, beyond compression

Besides providing the means to implement leading-edge compression technology, the standard provides the foundation for interoperable genomic information processing applications. ISO/IEC is also engaged in supporting the maintenance of the standard to guarantee the perenniality of the applications using MPEG-G technology. A list of the essential features of the MPEG-G technology is presented what follows.

### Selective access to compressed data

The indexing capabilities embedded in an MPEG-G file enable several types of selective access to the compressed data. Specifically, the following types of selective access, which can be combined in the same query, are supported:

- Genomic interval in terms of start to end mapping position on a given reference sequence
- Data type (i.e., a single Data Class)
- Sequence reads with number of substitutions below/above a certain threshold
- Sequence reads with multiple alignments
- Matching on previously defined patterns (e.g., barcodes) on raw or unmapped reads
- Labels on contiguous as well as non-contiguous intervals, possibly across multiple Datasets

### Data streaming

MPEG-G also provides the means for the efficient packetization of compressed data. This allows receiving devices to start processing the data before transmission is completed. The main capabilities of MPEG-G streaming are:

- Packet size adaptation to the channel characteristics/state
- On-the-fly indexing of streamed data
- Packet-based filtering of genomic data

Streaming within MPEG-G is enabled by the specification of a *Transport Format* which provides an extra set of data structures (packets and mapping tables, see Figure 3), in addition to those depicted in Figure 2 (which represents the File Format). The Transport Format allows multiplexing the File Format structures into data streams, of which each is composed of multiple packets that can be dynamically adapted to the network characteristics and conditions. Furthermore, the Transport Format allows for error detection, out-of-order delivery and re-transmission of erroneous/incomplete data on the protocol level (for example TCP/IP). File and Transport Formats are mutually convertible with no loss of information via a normative conversion process defined in the MPEG-G standard.

**Figure 3:**
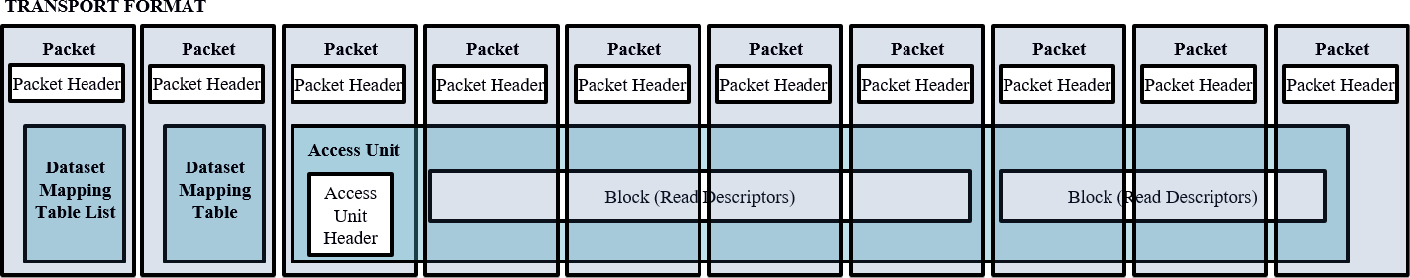
Key components of the MPEG-G Transport Format.

### Aggregation of genomic studies and incremental update of sequencing data and meta-data

Multiple related genomic studies can be encapsulated in the same MPEG-G file while still being separately accessible. This is supported by the notions of Datasets and Dataset Groups. A Dataset typically contains the result of a sequencing run, and a Dataset Group typically contains the runs associated to the same study. Aggregating (parts of) studies stored in multiple files is supported by a mechanism of file concatenation which does not require the re-coding of the compressed data. The same is true for a single Dataset that can be integrated with additional Access Units without the need to decompress and re-compress the existing Access Units.

For the aggregation, only the update of the indexing information and part of the associated metadata is required. Once the different studies have been aggregated, transversal queries over multiple studies are possible (e.g., “select chromosome 1 of all compressed samples”).

### Enforcement of privacy rules

Data encoded in an MPEG-G file can be linked to multiple owner-defined privacy rules, which impose restrictions on data access and usage. MPEG-G provides a syntax to express a hierarchy of privacy rules to be enforced on the coded content. This enables for example the implementation of the delegation of rights among different users. The data owner might delegate different levels of access permission to different users such that the personal physician will have higher access privileges than a research center performing a study on a large population.

### Selective encryption of sequencing data and metadata

The encryption of genomic information is supported by MPEG-G at different levels in the hierarchy of MPEG-G logical data structures. Each identifiable portion of the coded file can be associated to access control mechanisms such as encryption or digital signatures. The granularity of the protection mechanisms ranges from the encryption of a few features of aligned reads (e.g., mapping positions) to the entire Dataset or Dataset Group. MPEG-G does not enforce any specific selection of elements to encrypt, but provides a syntax to support any type of strategy. However, some parameters, such as the cipher, are restricted to sets of values in order to respect the security recommendations and simplify implementation and compatibility. This approach permits to limit the potentially resource-intensive application of data protection only to those portions of data which really need to be protected, leaving in clear text the non-sensible data.

### Annotation and linkage of genomic segments in the compressed domain

The MPEG-G specification provides a standard syntax to associate metadata to compressed genomic data for the implementation of annotation mechanisms. Additionally, MPEG-G provides support for linking segments within a single genomic sample or across multiple genomic samples. To this end, MPEG-G supports the aggregation of different blocks of compressed genomic data so that retrieval can be performed with a single query. The mechanism relies on the notion of associating a textual identifier to a syntax expressing the characteristics of the genomic data to be aggregated. Such characteristics can be the genomic interval on a reference sequence, the type of data (i.e., Data Class), or the Dataset identifier. This enables linking genomic regions that can be far away from each other (e.g., on different chromosomes or from different sequencing runs) and as such simplifies the annotation and retrieval of data.

### Interoperability with main existing technologies and formats

Conversion to/from formats such as FASTQ, SAM or BAM is supported by MPEG-G. The MPEG-G specification provides guidelines on how to transcode existing content to MPEG-G and back to its original format. This is particularly useful especially for those cases where the transcoding mechanism cannot be inferred unambiguously from the SAM specification.

### MPEG-G implementation framework

The MPEG-G specifications comprise normative *Reference Software* and *Conformance* testing, which complement the formal specifications of syntax, semantics, and decoding process with tools and methodology for robust and reliable Conformance validation. Besides, a Genomic Information Database was also compiled during the development of the standard. This database contains a collection of sequencing data used to assess the performance of genomic information compression technologies.

### Reference Software

To support and guide the implementation of MPEG-G, the standard includes a normative Reference Software. The Reference Software is normative in the sense that any conforming implementation of the decoder, taking the same conformant compressed bitstreams, and using the same normative output data structures, will output the same data. That said, complying MPEG-G implementations are not expected to follow the algorithms, or even the programming techniques used by the Reference Software; such software is solely intended as a support to the process of developing implementations of an ecosystem of compliant devices and applications. Hence, the availability of a normative implementation is only an additional support to the textual specification. It should also be underlined that the Reference Software is not intended as an optimized implementation of an MPEG-G decoder. This also means that the Reference Software should not be used as a benchmark of performance.

### Conformance

Conformance is fundamental in providing means to test and validate the correct implementation of the MPEG-G technology in different devices and applications, and to ensure interoperability among all systems. Conformance testing specifies a normative procedure to assess conformity to the standard on an exhaustive set of compressed data: every decoder claiming MPEG-G conformance will have to demonstrate the correct decoding of the complete conformance testbed.

The set of bitstreams for Conformance testing is available in the MPEG-G Genomic Information Database (see below).

### MPEG-G Genomic Information Database

The MPEG-G Genomic Information Database is a collection of statistically meaningful sequencing data used to assess the performance of genomic information compression technologies. Besides the actual sequencing data, the database contains a set of reference sequences and supporting data needed for variant calling experiments (see Methods). When compiling the database special emphasis was put on incorporating data with as much diversity as possible. Hence, it contains data generated by different sequencing technologies, produced for the purpose of conducting different experiment types (e.g., WGS, RNA-seq, etc.), and originating from samples across different species such as *H. sapiens*, *D. melanogaster* or *E. coli*.

## DISCUSSION

Widely used formats to represent genomic information (i.e., FASTQ and SAM/BAM) were designed when sequencing data was scarce and precious, and the range of applications limited. Typical shortcomings include under-specified application programming interfaces preventing the creation of inter-operabile applications and devices; no framework to enforce privacy protection; undocumented process for extensions and amendments; poor compression performance (in some cases limited to generalized compressors applied to plain text); lack of conformance tests; no transport format specified, nor support for packetized data streaming. The new paradigm introduced by high-throughput sequencing machines—relatively inexpensive high coverage sequencing, with an almost infinite number of derived biological protocols and downstream analysis workflows— strongly encourages the adoption of a more sophisticated way to store, handle and share genomic data. Solutions able to process sequencing data with high efficiency, similar to those available today to share digital media content, will only be possible if all the fundamental design problems present in the existing formats are identified and solved. MPEG-G represents an important step in that direction. It paves the way to novel software solutions that will allow independent groups and organizations around the world to seamlessly communicate and share data, without losing inter-operability with existing applications.

The analogy to the digital media industry can be carried on further. MPEG-G aims to make genomic data access, processing and sharing—either in the cloud or on local storage—as simple as streaming an audio file or watching a movie. In other words, the MPEG-G specification will hopefully enable for genomics the same ground-breaking developments that the digital media industry has witnessed between the end of the past century and the beginning of the current one. One of the main drivers of that revolution was the impressive performance of data compressors that enabled digital media storage and transfer on a scale never experienced before. Another determinant has been the open and fair process of technology evaluation and specification under the supervision of international and neutral institutions such as ISO and IEC. This encouraged small and large organizations around the world to join forces and work in a collaborative environment, at the same time providing reassurance on the long-term stability of the standard, which is a public document maintained by a large group of experts. Finally, the specification of standard interfaces for systems interoperability enabled the proliferation of compatible technology and products which make up the digital media ecosystem as we know it today. These very elements are present in MPEG-G; hopefully they will facilitate the creation of a similar ecosystem (see Figure 4) that will eventually democratize genomic applications such as personalized medicine.

**Figure 4:**
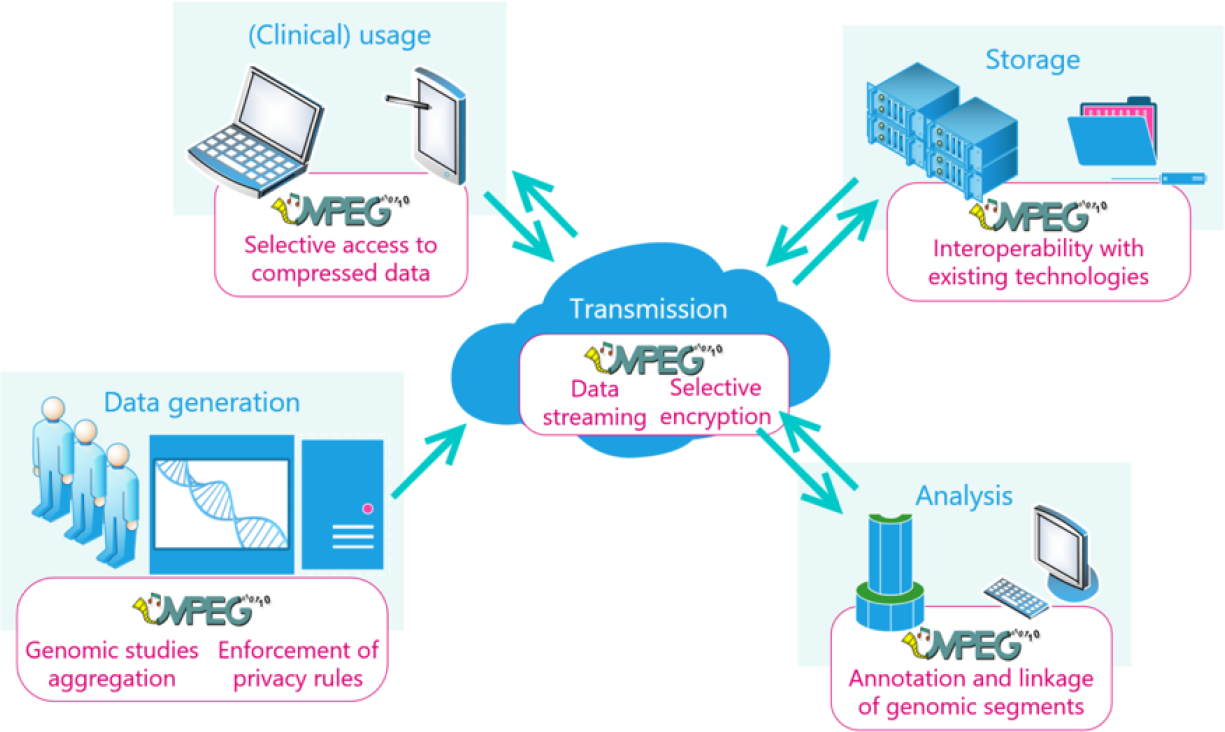
A genomic ecosystem fueled by MPEG-G.

With the release of the MPEG-G standard, genomic data sharing will benefit from both the reduced file sizes and the standardized interfaces; analysis tools will be offered with sophisticated ways to access and manipulate data; security and privacy protection will be seamlessly implemented thanks to built-in access control. Applications which are today conceivable but not easily implemented due to either the sheer amount of data to be transferred or prohibitive IT development costs may soon become a reality. In addition, the open transparent process of maintenance and update of the standard will encourage institutions to invest in the adoption and further development of the technology. Hence, MPEG-G will hopefully continue to evolve, improve and bear fruit for a long time in the future.

## METHODS

In this section we more formally describe the MPEG-G standard and its components. In particular, the MPEG-G standard is divided into the following five parts.

<u>Part 1</u>: Transport and Storage of Genomic Information. This part of the standard specifies how the genomic data is organized within MPEG-G structures for transport (i.e., streaming) and storage. A reference conversion process is provided to go from the File Format to the Transport Format and vice versa.

<u>Part 2</u>: Coding of Genomic Information. This part specifies the syntax used to represent unaligned (e.g., raw) and aligned sequence reads and the associated identifiers, quality values and reference sequences, if any. This is the part of the standard that deals with compression by describing the normative behavior of a compliant decoder parsing an MPEG-G bitstream. Only the decoding process is specified while any encoding algorithm can be used, providing it produces a bitstream compliant with this part of the standard.

<u>Part 3</u>: Metadata and APIs. This part of the standard specifies how an MPEG-G compliant bitstream can be integrated with metadata describing, for example, a genomic study or a sequencing run. Other topics covered by this part include the specification of normative interfaces to access MPEG-G data from external systems, the specification of mechanisms to implement access control, integrity verification, as well as authentication and authorization mechanisms. This part also includes an informative section devoted to the mapping between SAM and MPEG-G data structures.

<u>Part 4:</u> Reference Software. To support and guide potential implementers of MPEG-G, the standard includes a normative Reference Software. The Reference Software is normative in the sense that any conforming implementation of the decoder, taking the same conformant compressed bitstreams and using the same normative output data structures, will output the same data.

<u>Part 5</u>: Conformance. This part of the standard is fundamental in providing means to test and validate the correct implementation of the MPEG-G technology in different devices and applications to ensure the interoperability among all systems. Conformance testing specifies a normative procedure to assess conformity to the standard on an exhaustive set of compressed data.

A more detailed description of Part 1, 2 and 3 is provided in what follows.

### Part 1: Transport and Storage of Genomic Information

MPEG-G specifies a digital container format for transmission and storage of the genomic data compressed according to Part 2 of the standard. In MPEG jargon the container format used for the transport of packetized data (i.e., stream) on a telecommunication network is referred to as *Transport Format*, while the container format used for storage on a physical medium (i.e., file) is referred to as *File Format*. The process of converting a stream to a file and vice versa is normative and specified in the standard.

#### File Format

An MPEG-G file is organized in a file header and one or more containers named Dataset Groups. Each Dataset Group contains a Dataset Group header, optional metadata containers and encapsulates one or more Datasets. Each Dataset has a Dataset Header, optional metadata containers and carries one or more Access Units. The Access Unit is the actual container of the compressed genomic data. It includes an Access Unit header which provides a description of the compressed content (type of data, number of reads, genomic region including the compressed reads, etc.).

In the case where an MPEG-G file was constructed without Descriptor Streams, the Access Unit contains a collection of *Blocks* of coded information that can be decoded independently using global data at the Dataset level and eventually information contained in other Access Units, such as Access Units containing data of an MPEG-G encoded reference sequence. Otherwise, the blocks of coded information are grouped by type, concatenated and stored as Descriptor Streams. The index mechanism then allows to associate a given Access Unit to the corresponding collection of Blocks. In either case, each Block is compressed using the entropy coding techniques (see Part 2 below) most suitable to the measured statistical properties. This nested data structure is depicted in Figure 2.

#### Transport Format

In addition to the data containers defined for the File Format, MPEG-G specifies data structures supporting packetized data transport over a network. Such structures are defined both to carry the compressed genomic data and to update metadata describing the streamed content. An example of the latter type of data is indexing information used by the receiving end to enable selective access even on partially transmitted content.

The Transport Format structures are instrumental in the specification of the normative process of conversion between Transport Format (i.e., an MPEG-G stream transmitted over the internet) and File Format (i.e., an MPEG-G file stored on disk).

### Part 2: Coding of Genomic Information

Genomic Records are classified into six Data Classes according to the result of the primary alignment(s) of their reads against one or more reference sequences as shown in Table 1. Records are classified according to the types of mismatches with respect to the reference sequences used for alignment.

**Table 1:** Data Classes defined in MPEG-G.

| Class name | Semantics |
| --- | --- |
| P | Reads perfectly matching to the reference sequence. |
| N | Reads containing mismatches which are unknown bases only. |
| M | Reads containing at least one substitution, and possibly unknown bases, but no insertions, no deletions and no clipped bases. |
| I | Reads containing at least one insertion, deletion or clipped base, and possibly unknown bases or substitutions. |
| HM | Half-mapped pairs where only one read is mapped. |
| U | Unmapped reads. |

To further improve compression efficiency, the information contained in the clustered Genomic Records is split across Descriptor Streams. The concept of splitting the information contained in the clustered Genomic Records into Descriptor Streams allows tailoring encoding parameters according to the statistical properties of each Descriptor Stream [3].

#### Compression modes for raw sequencing data

Raw sequencing data can be encoded according to two different approaches, depending on the application at hand:

##### High compression ratio and indexing

A high compression ratio is reached by leveraging the high redundancy in genomic sequence data. This enables the use of well-known compression techniques such as differential coding of sequences with respect to already encoded data. This approach achieves a maximum compression ratio but requires the availability of the entire Dataset as well as a few preprocessing stages which may impact the compression latency. Differential coding of raw genomic sequences relies on the identification of common patterns (i.e., “signatures”) shared among several sequences. These common patterns are encoded only once along with the nucleotides specific to each read (i.e., the “residuals”). The presence of such signatures enables the implementation of indexing schemes with which the compressed data can be searched by means of pattern matching algorithms. This mode is suitable, for example, for long term storage of raw sequencing data. Examples of preprocessing technologies of which the data output can be represented using the decoder syntax defined in MPEG-G for this mode of operation include those presented in ORCOM [13], HARC [14], FaStore [7] and, in general, all (future) preprocessing technologies that cluster reads based on common patterns.

##### Low latency

When low streaming latency has higher priority than compression ratio, MPEG-G also supports a “high throughput” compression approach which can be applied as soon as genomic sequences become available. In such case no data preprocessing over the entire Dataset is required prior to the actual encoding. This approach enables streaming scenarios in which the genomic data need to be transmitted to a remote device from the sequencing facility as soon as they are available (and possibly even before the sequencing process has been completed).

#### Compression modes for aligned reads

Genomic sequence reads mapped onto reference sequences can be compressed following two approaches:

##### Reference-based compression

In this approach the genomic sequences are represented by the differences they present with respect to the reference sequences as well as by the associated alignment information. The reference sequences can be embedded as Datasets within the same MPEG-G file. Optionally, external reference sequences can be used. MPEG-G specifies how external reference sequences can be identified unambiguously using a URI, checksums, etc. External reference sequences can either be delivered to a decoder as MPEG-G Datasets or in FASTA format.

##### Reference-free compression

In this approach the aligned sequences are compressed without referring to any reference sequence. A local assembly of the underlying sequence is built per group of reads, and reference-based compression with respect to the computed local assembly is then applied. In this case, there is no need to have access to any reference sequences neither at the encoder nor at the decoder side. The two main technologies implementing this approach and submitted for consideration include TSC [15] and DeeZ [8].

#### Compression modes for quality values

Due to their higher entropy and larger alphabet, quality values have proven more difficult to compress than the reads [16, 11]. In addition, there is evidence that quality values are inherently noisy, and downstream applications that use them do so in varying heuristic manners. As a result, quantization of quality values can not only significantly alleviate storage requirements but also provide variant-calling performance comparable–and sometimes superior–to the performance achieved using the uncompressed data (see [16, 11] and references therein). This is possible with even 0.5 bits per quality score rather than the 3 bits needed for lossless compression, approximately. Therefore, in MPEG-G, quality values can be encoded either in a lossless manner or in a quantized manner. When encoding quality values losslessly, several transformations can be applied to the quality values prior to the actual arithmetic coding (see also step 2 of the entropy coding procedure below). These transformations include, among others, differential coding, run-length encoding, and a transformation named “match coding” which can be regarded as a modified Lempel-Ziv scheme [17].

Quantization of quality values, however, can lead to a dramatic reduction of the bitstream size after entropy coding. However, to facilitate minimization of any quantization effects, the MPEG-G standard provides several mechanisms to allow an encoder to perform a fine-grained selection of quantization schemes.

In the case of unaligned reads, an MPEG-G compliant encoder is free to choose any beneficial quantization scheme. This includes quantization schemes of recently published research such as [18, 19, 20, 9]. The specific quantization scheme used is signaled to a decoder by the means of a *Quality Value Codebook*. The quantized quality values are signaled to a decoder as *Quality Value Indexes* into this Quality Value Codebook.

In the case of aligned reads, MPEG-G introduces an additional dimension to fine-tune quality value quantization: codebooks can be chosen per genomic position, i.e., per locus [10]. Therefore, one Quality Value Codebook Identifier per genomic position is sent to a decoder along with the Quality Value Indexes.

In both cases, before entropy coding, the Quality Value Indexes are split into separate streams per Quality Value Codebook. Finally, an MPEG-G encoder is also allowed to tune the quantization by selecting different codebooks per Data Class as well as per Access Unit.

#### Compression modes for read identifiers

Read identifiers are broken down into a series of tokens which can be of three main types: strings, digits and single characters. A read identifier is represented as a set of differences and matches with respect to one of the previously decoded Read Identifiers. This approach does not rely on any sequencing manufacturer implementation and only assumes that within the same sequencing run the structure of read identifiers is mostly constant. Note that this method (or variants of it) for identifiers compression has been previously employed in compressors such as SamComp [21], Quip [22], DeeZ [8], and FaStore [7], among others.

#### Compression modes for reference sequences

MPEG-G supports the use of reference sequences both in the FASTA format and in the MPEG-G compressed format. The reference sequences can also be embedded as Datasets within the same MPEG-G file. Optionally, external reference sequences (i.e., sequences that are not included in the bitstream) can be used. MPEG-G specifies how external reference sequences can be identified unambiguously using a URI, checksums, etc.

A reference sequence in MPEG-G can be coded either as a stand-alone Dataset or as a difference with respect to another reference sequence. In the first case the reference sequence is coded as a sequence of Genomic Records belonging to Data Class U (unmapped reads) and encoded with the approaches described for unmapped sequence data. In case of differential encoding with respect to another reference sequence, the same approaches used for aligned data are used. In this case Genomic Records belonging to Data Classes P, N, M, I and U can be used to represent segments of the encoded assembly. In case of differential coding, the reference sequence used as reference to code one or more other genome sequences does not need to be a real genome sequence but can be synthesized to improve the compression performance. This can be helpful when compressing collections of genome sequences using a common reference which is not necessarily one of the sequences of the collection.

#### Entropy coding

Storing different types of data in separate Descriptor Streams allows for a significantly higher compression effectiveness. The different statistical properties of each descriptor can be exploited to define different source models to be used for entropy coding. The increased compression efficiency is generated by the adoption of the appropriate context adaptive probability models according to the statistical properties of each source model.

To compress the heterogeneous set of descriptors, MPEG-G specifies the use of Context-Adaptive Binary Arithmetic Coding (CABAC) [23], as used in popular video coding standards and the genomic data compression solutions AFRESh and AQUa [24, 25]. By selecting this highly-effective arithmetic coder, the implementation of compliant codecs is simplified significantly, as a wide range of implementations, both in hardware and in software, are currently available.

The compression process consists of 5 steps (see Figure 5): input data parsing, value transformation, value binarization, context selection, and CABAC.

**Figure 5:**
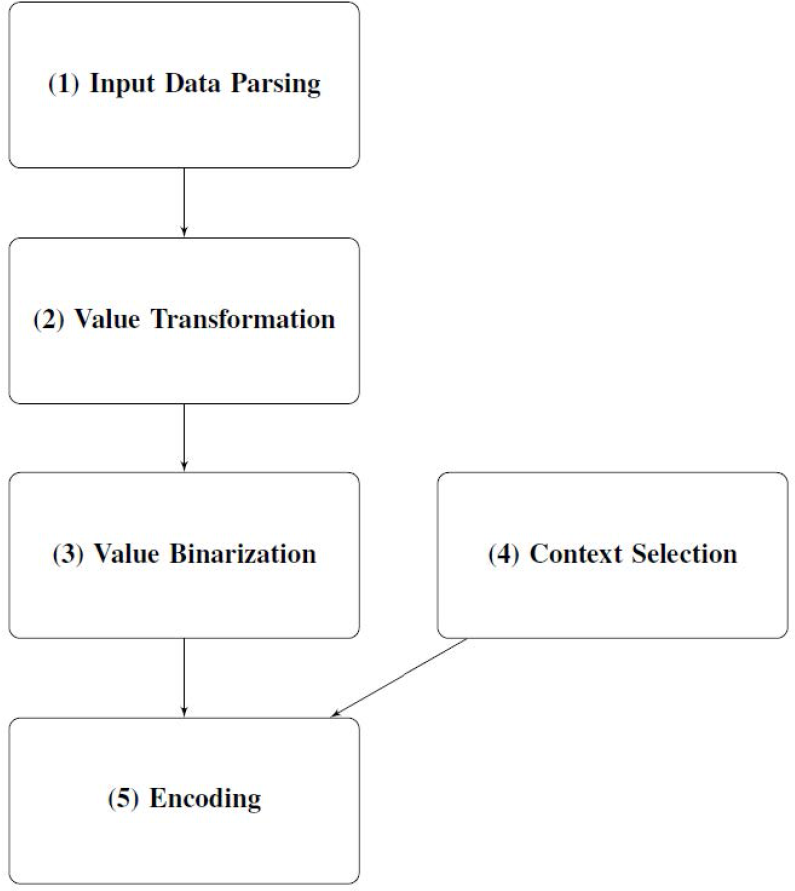
Use of context-adaptive entropy coding in MPEG-G.

In step 1 (input data parsing) the Descriptor Streams are parsed as a sequence of values. If beneficial, the data contained in a Descriptor Stream can be split into multiple subsequences. Each of the resulting subsequence is processed separately in steps 2 to 5.

In step 2 (value transformation) an optional (sequence of) transformation(s) is applied to the values produced by step 1. Some of the transformations generate additional data streams. Each of the resulting transformed subsequences is processed separately in steps 3 to 5.

In step 3 (value binarization) the values in the different transformed subsequences are converted to a binarized representation (i.e., a set of bits). To allow for effective compression in step 5, the combination of transformation and binarization should be selected in such a way that the value of each bit of the binarization is as predictable as possible. The stream of binarizations serves as input for step 5.

In step 4 (context selection) the context sets that will be used during the encoding step (step 5) are identified. Each context set contains the contexts required to encode one input value. The goal of the context selection step is to select the context set that is expected to resemble as much as possible the distribution of the bits in the binarized representations generated at step 3. The stream of selected contexts serves as support values for the entropy encoder in step 5.

In step 5 (CABAC) the bins of the binarized representations generated at step 3 are encoded, using the context sets that have been selected in step 4 or in bypass mode, using a non-adaptive context that represents equiprobability.

#### Decoding process

The MPEG-G specification not only defines the syntax and semantics of the compressed genome sequencing data, but also the deterministic decoding process.

The normative input of an MPEG-G decoding process is a concatenation of data structures called *Data Units*. Data Units can be of three types according to the type of conveyed data. A

Data Unit of type 0 encapsulates the decoded representation of one or more reference sequences, a Data Unit of type 1 contains parameters used during the decoding process in a structure called Parameter Set, and a Data Unit of type 2 contains one Access Unit.

Data Units of type 0 and 1 are used during the decoding process of Data Units of type 2 but do not produce any normative output. The data carried by such Data Units are managed by the decoding process in an implementation-dependent way. It is the decoding process of Access Units that produces a normative output either in the form of MPEG-G Records for Access Units containing raw or aligned reads or in the form of a Raw Reference structure for Access Units containing a compressed reference sequence or a part thereof. An MPEG-G Record can be regarded as an improved SAM record: in MPEG-G read pairs are typically coded in the same record unless certain conditions are met, such as the pairing distance is above a user-defined threshold, or the mate is mapped to a different reference sequence. The decision to split a read pair is taken by the encoder and the pairing information is transmitted to the decoder for each read in the pair using the appropriate descriptors. The decoding process is fully specified such that all decoders that conform to Part 2 of the standard will produce identical decoded outputs. The normative decoding process includes all hierarchies of data structures, from the multiplexed bitstreams included in MPEG-G files or the data streams in streaming scenarios, to the descriptors Blocks and to the normative output. A simplified diagram of the decoding process is shown in Figure 6.

**Figure 6:**
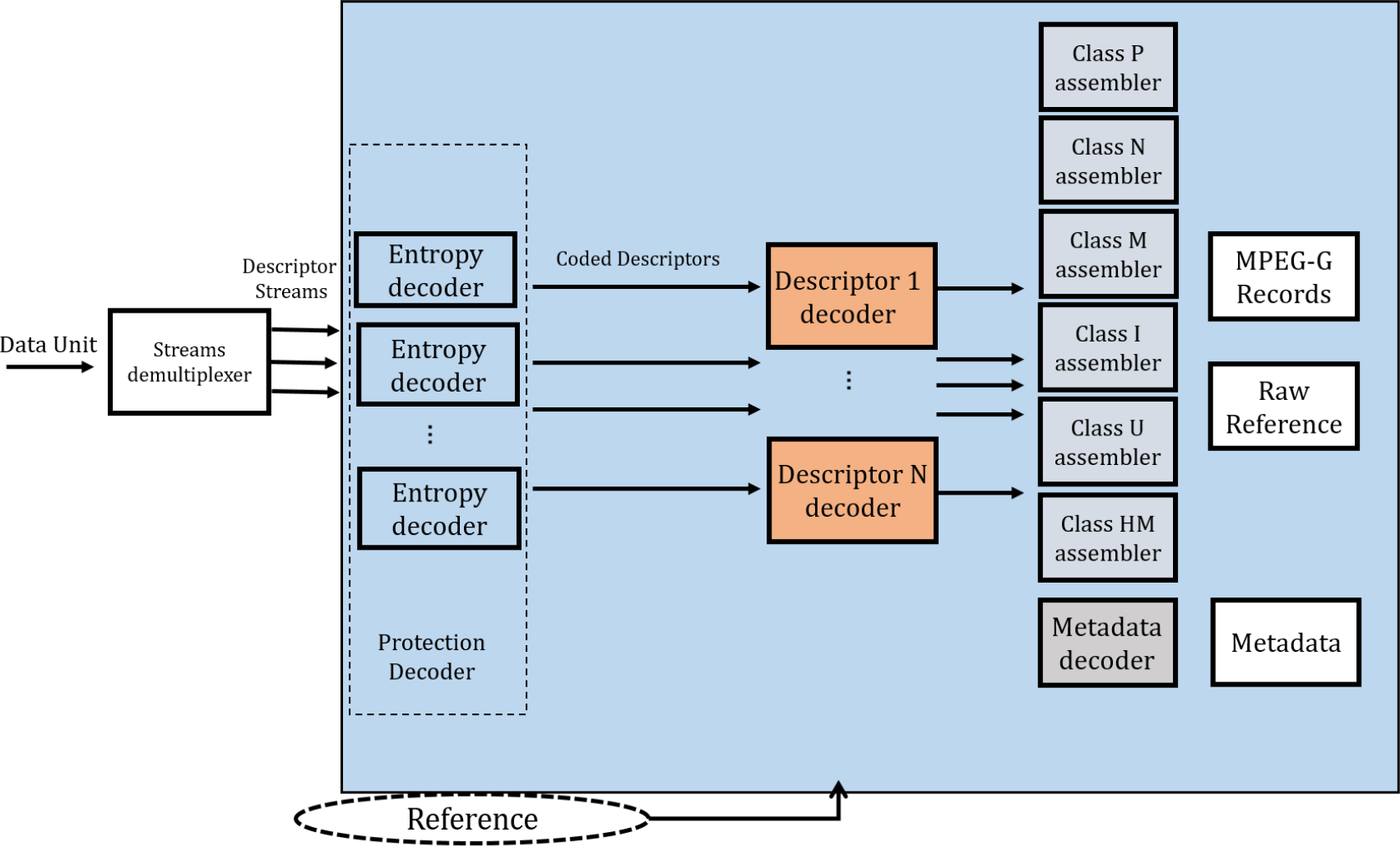
Simplified MPEG-G decoding process from a Data Unit to the normative output.

### Part 3: Metadata and APIs

#### Enforcement of privacy rules

Data encoded in an MPEG-G file can be linked to multiple owner-defined privacy rules, which impose restrictions on data access and usage. The privacy rules are specified in XACML (eXtensible Access Control Markup Language) Version 3.0, an OASIS standard [26].

The MPEG-G file format includes protection information provided in specific containers available at most levels of the MPEG-G hierarchy, including Dataset Group, Dataset, Descriptor Stream and Access Unit levels. These protection containers provide—in addition to the privacy rules to be applied to the information they refer to—mechanisms to manage the confidentiality and integrity of the information. Specifically, the information on privacy rules is only available at the Dataset Group and Dataset levels. A privacy rule could specify, for example, access control to specific regions related to identifying Alzheimer predisposition. By using encryption techniques combined with privacy rules the genomic data is efficiently protected from unauthorized access. Therefore, only users authorized by the rules can perform operations over protected regions.

#### Encryption of sequencing data and metadata

MPEG-G supports the encryption of genomic information at different levels in its hierarchy of logical data structures. The protection information specifies how the data structures at the same level, and the protection information containers of a layer immediately below, are encrypted. This information is represented using the XML Encryption v1.1 standard (http://www.w3.org/TR/xmlenc-core1). On the other hand, authentication and integrity may be provided by means of electronic signatures using XML Signature v1.1 (http://www.w3.org/TR/xmldsig-core1).

Encryption is not only possible for the “low-level” detailed sequencing data and metadata included in Genomic Records, but also for the “high-level” metadata available for the Dataset Group and Dataset hierarchy levels. For this purpose, MPEG-G provides metadata information structures specified using XML v1.1 (http://www.w3.org/TR/xml11), with a set of elements for those levels. It includes a minimum core set of metadata elements (such as title and samples for Dataset Groups, or title and project centres for Datasets). Users and applications can extend this core set, in a standardized way, by including extra information elements.

In addition, metadata profiles are specific subsets of metadata sets specified using mechanisms also provided in the standard. A specified metadata profile may correspond to a common metadata set specified or used out of MPEG-G, such as those from the European Genome-phenome Archive (EGA) and the National Cancer Institute (NCI) Genomic Data Commons (GDC). A metadata profile includes a subset of core elements and a set of new elements specified with the extensions mechanism.

#### The MPEG-G Genomic Information Database

The sequencing data contained in the database is classified according to: i) experiment type, ii) sequenced organism, and iii) employed sequencing technology. The database includes data related to WGS, metagenomics sequencing, RNA sequencing, and cancer sequencing experiments. The WGS data is further extended with simulated human WGS data. Furthermore, data from a wide variety of taxa—namely Animalia, Plantae, Fungi, Bacteria and Viruses—is included in the database: *Drosophila melanogaster* and *Homo sapiens* (Animalia), *Theobroma cacao* (Plantae), *Saccharomyces cerevisiae* (Fungi), different strains of *Escherichia coli* and *Pseudomonas aeruginosa* (Bacteria), and ΦX174 (Viruses). Finally, the data was produced using different sequencing technologies: i) sequencing by synthesis (Illumina/Solexa Genome Analyzer, Illumina Genome An-alyzer IIx, Illumina MiSeq, Illumina HiSeq 2000, Illumina HiSeq X Ten, Illumina NovaSeq 6000); ii) single molecule real time sequencing (Pacific Biosciences SMRT (PacBio)); iii) nanopore sequencing (Oxford Nanopore MinION); and iv) ion semiconductor sequencing (Ion Torrent PGM).

The database can be accessed at https://github.com/voges/mpeg-g-gidb

#### MPEG-G documents

Due to the size and breadth of the MPEG-G standard we refer the reader to the official documents that describe it. These documents will be publicly available when the final version is published by ISO and IEC. Intermediate public documents can be found on the MPEG website section devoted to MPEG-G (https://mpeg.chiariglione.org/standards/mpeg-g) and on the MPEG-G portal at https://mpeg-g.org.

## ACKNOWLEDGMENTS

This project was partially supported by the grant numbers 2018-182798 and 2018-182799 from the Chan Zuckerberg Initiative DAF, an advised fund SVCF; an SRI grant from UIUC; the Swiss Commission for Technology and Innovation (CTI), grant 19318.1; and the Spanish Government (GenCom, TEC2015-67774-C2-1-R).

